# Adipocyte-specific deletion of the oxygen-sensor PHD2 sustains elevated energy expenditure at thermoneutrality

**DOI:** 10.1101/2021.01.05.425401

**Authors:** Mario Gomez Salazar, Iris Pruñonosa Cervera, Rongling Wang, Karen French, Ruben García-Martín, Matthias Blüher, Christopher J Schofield, Roland H Stimson, Triantafyllos Chavakis, Elias F Gudmundsson, Lori L Jennings, Vilmundur G Gudnason, Nicholas M Morton, Valur Emilsson, Zoi Michailidou

## Abstract

Enhancing brown adipose tissue (BAT) function to combat metabolic disease is a promising therapeutic strategy. A major obstacle to this strategy is that a thermoneutral environment, relevant to most modern human living conditions, deactivates functional BAT. We showed that we can overcome the dormancy of BAT at thermoneutrality by inhibiting the main oxygen sensor HIF-prolyl hydroxylase, PHD2, specifically in adipocytes. Mice lacking adipocyte PHD2 (*P2*KO^ad^) and housed at thermoneutrality maintained greater BAT mass, had detectable UCP1 protein expression in BAT and higher energy expenditure. Mouse brown adipocytes treated with the pan-PHD inhibitor, FG2216, exhibited higher *Ucp1* mRNA and protein levels, effects that were abolished by antagonising the canonical PHD2 substrate, HIF-2a. Induction of *UCP1* mRNA expression by FG2216, was also confirmed in human adipocytes isolated from obese individuals. Human serum proteomics analysis of 5457 participants in the deeply phenotyped Age, Gene and Environment Study revealed that serum PHD2 (aka EGLN1) associates with increased risk of metabolic disease. Our data suggest adipose–selective PHD2 inhibition as a novel therapeutic strategy for metabolic disease and identify serum PHD2 as a potential biomarker.

## Introduction

Several elegant studies have identified approaches to activate thermogenic brown adipose tissue (BAT) and enhance metabolism (1–3). However, the majority of these experiments were conducted on rodents at ambient temperature (19–21°C), a condition where BAT is already active (4–6). Consequently, translation of these studies is questionable, because humans predominantly live in thermoneutral conditions by using clothing or other means to maintain thermal homeostasis (7). One highly upregulated gene in adipose tissue during cold exposure of rodents is the hypoxia inducible transcription factor (HIF)-2α (8), suggesting that the hypoxia signalling pathway is involved in BAT activation. The oxygen-sensing HIF-prolyl hydroxylases (PHDs) are central regulators of the hypoxic response (9–10). PHD2 is the dominant oxygen sensor and is a negative regulator of HIF-a activity in normoxia (9–10). PHD2 hydroxylates key proline residues on HIF-a isoforms in normoxia, leading to its proteasomal degradation (9–10).

PHD inhibitors are of increasingly medical significance and have already advanced to phase 3 clinical trials for the treatment of chronic kidney disease (CKD) (11) and recently been licenced for clinical use in China and Japan (12–13). Previously, we and others, have shown that models of whole-body or adipose-specific deletion of the dominant ubiquitously expressed PHD2 isoform induced protection from metabolic disease, in part by reducing plasma lipid levels (14–15). We hypothesized that the lipid-lowering effects of PHD2 deficiency is due to enhanced BAT thermogenesis. Here we report on the testing of this hypothesis, using a genetic mouse model of adipose tissue–specific deletion of PHD2 (*P2*KO^ad^) under thermoneutrality, to better align with human metabolism, and thus enhance the translational relevance of our findings. To address the relevance of PHD2 in human metabolic disease traits, we interrogated the unique large–scale serum proteomics and deep metabolic and anthropometric phenotyping data from the Age/Gene Environment Susceptibility study (Icelandic Heart Association) (16–17).

## Results and Discussion

### Loss of adipocyte PHD2 mediates retention of BAT function at thermoneutrality

Housing of mice at thermoneutrality (TN; approximately 28-30°C) results in loss of BAT function (18–19). This aligns rodent metabolism more closely with that of humans, providing a better preclinical modelling system (5–7, 18–20). We tested whether the metabolic advantage observed with adipose-specific PHD2 deletion previously (15,21), was retained at TN (28°C), thus indicating an effect independent from, or maintenance of BAT function.

Adiponectin–*Cre* (22) mice were crossed with *PHD2^fl/fl^* mice (15) to delete PHD2 (referred as *P2*KO^ad^) in white (data not shown) and brown adipocytes (Supplemental Fig. 1a). This approach targets all white (WAT), beige and BAT adipocytes (23) and thus reflects a generalised adipose deletion not restricted only to the rodent-specific BAT depot. *P2*KO^ad^ mice had >85% reduction in *Phd2* mRNA levels compared to control littermates, and consequently elevated levels (1.6-fold, p=0.018) of the classical HIF-target gene *Vegfa* in BAT (Supplemental Fig. 1a) and in inguinal white adipose tissue (iWAT, mean±SEM: Control: 0.38±0.07 *vs P2*KO^ad^: 0.73±0.12, p=0.045). This was associated with HIF-1a stabilisation, as expected (Supplemental Fig. 1b), irrespective of oxygen levels in the tissue. This data confirms enhanced vascularization in the adipose tissues in this model as we previously showed in WAT irrespective of temperature (15). Unexpectedly, we found that despite similar body weights (Figure 1a), *P2*KO^ad^ mice had relatively higher energy expenditure (EE), especially in the dark phase (Figure 1b) than control littermates, without any significant changes in their activity levels (Figure 1c). However, *P2*KO^ad^ had a higher food intake (Figure 1d) compared to their littermate controls, consistent with balance of their higher EE. At thermoneutrality, body composition was similar between the genotypes (lean and fat mass, Figure 1e). However, the control littermate mice lost BAT mass (Supplemental Fig.1c) at TN, as expected, whereas the *P2KO^ad^* mice, maintained BAT mass (Supplemental Fig.1c) and had significantly greater BAT tissue mass (Figure 1f). White adipose depots (Figure 1g) and liver mass (Supplemental Fig. 1d) were comparable between genotypes.

**Figure 1.**
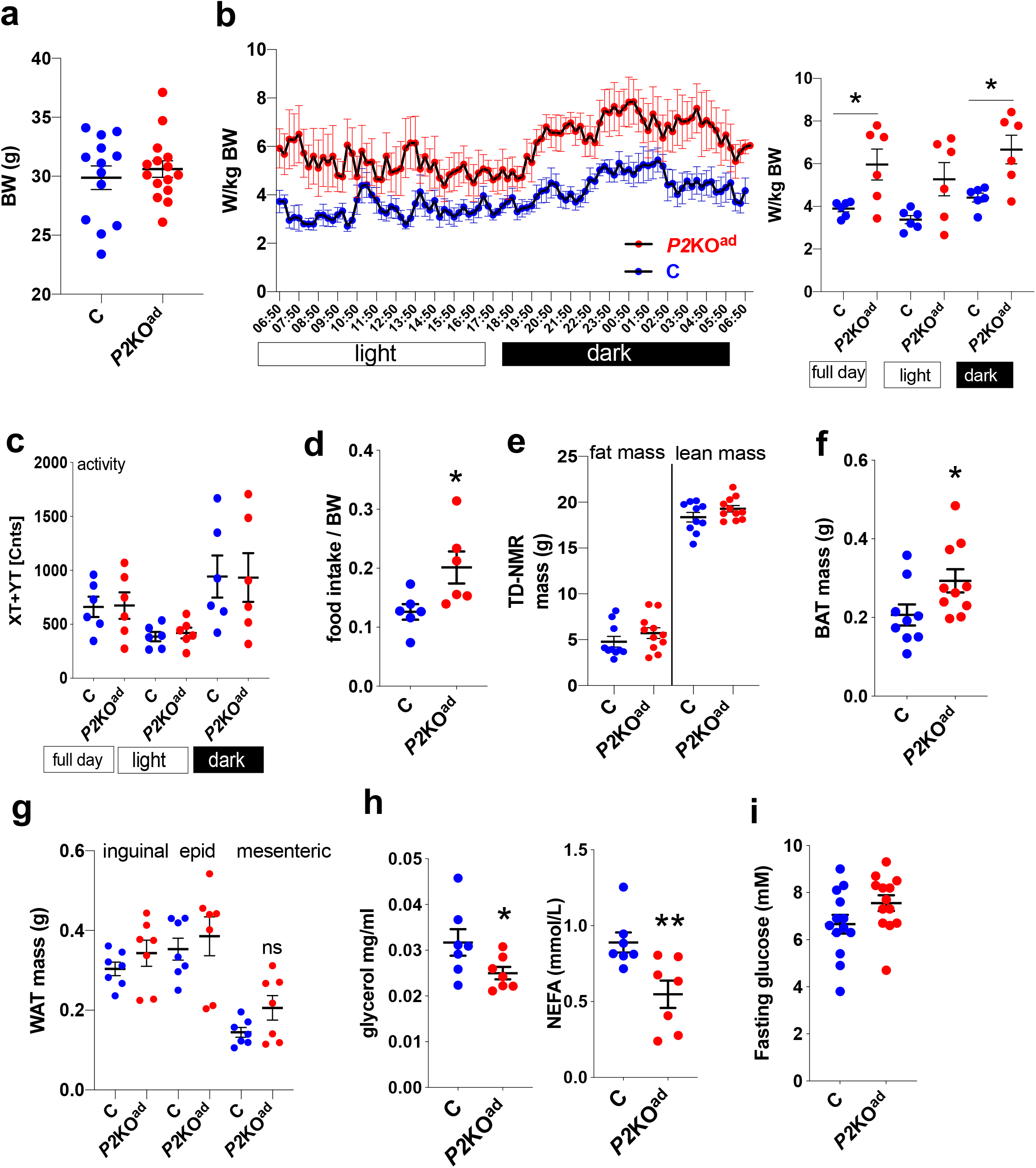
Adipose *Phd2* deletion permits BAT mass maintenance at TN. P2KO^ad^ (red) mice housed at 28°C for 3days have similar body weight **(a)** to control (blue) littermates but higher energy expenditure **(b)** (n=6-7), similar activity levels **(c)** and eat more **(d)**. Despite similar fat and lean mass measured by time-domain (TD) NMR (n=10-11) **(e)**, P2KO^ad^ mice have bigger BAT mass **(f)**, but a similar WAT mass **(g)**. P2KO^ad^ mice have lower plasma glycerol and non-esterified fatty acids (NEFA) (n=7) **(h)** but similar fasting blood glucose (n=13) levels **(i)**. Data are presented as mean+/− SEM. *p<0.05, ** p<0.01 by Student t-test.

Metabolically, during TN, the main effect of *adipose-Phd2* deletion was on plasma lipids (Figure 1h), as glucose levels were unaffected (Figure 1i). Specifically, *P2*KO^ad^ mice had significantly lower circulating glycerol and non-esterified fatty acids (NEFA) levels compared to control mice (Figure 1h). Because the main source of circulating NEFA and glycerol is white adipose tissue lipolysis, we measured NEFA and glycerol in inguinal white adipose (iWAT) explants in mice housed at TN. In accordance with the plasma levels, iWAT explants from *P2*KO^ad^ mice released less NEFA and glycerol in the culture media under basal conditions (Supplemental Fig.1e).

### Loss of Phd2 in adipocytes increases brown adipocyte proliferation

The main characteristic during BAT activation in cold exposure or beta-adrenergic stimulation is BAT hyperplasia and/or hypertrophy (24–25). To investigate the cellular basis for the unexpected resistance of BAT to regress at TN in *P2*KO^ad^ mice, we performed histological analysis. *P2*KO^ad^ mice had bigger adipocytes, in the range of 450-1000μm^2^, and more total brown adipocyte numbers (Figure 2a-b) compared to control mice, suggesting both hypertrophy and BAT hyperplasia are involved. More BAT does not necessarily mean functional BAT, we therefore, quantified UCP1+ cells at thermoneutrality using antibody staining as a proxy for functional BAT. *P2*KO^ad^ mice exhibited more UCP1 expressing cells (Figure 2c-d, Supplemental Figure 2a) in BAT than control littermates and more ki67+ cells (Figure 2c-d, Supplemental Fig.2b), suggesting higher proliferation rates. Deletion of adipocyte *Phd2* also led to higher levels of the BAT-activating adrenergic receptor, *Adrb3,* and *Ucp1* mRNA levels in BAT (Supplemental Fig.1f) and in iWAT (Supplemental Fig.1g). Previously we have shown that deletion of *Phd2* leads to better WAT vascularization (increased *Vegfa* mRNA/CD31 immunoreactivity) (15) at ambient housing conditions (21 °C). We confirmed here that *Vegfa* mRNA levels remain higher at thermoneutrality in BAT from *P2*KO^ad^ mice (Supplemental Fig.1a). Immunofluorescence for isolectin IB4, a vessel marker, also confirmed more extensive vascularization of BAT from *P2*KO^ad^ mice (Figure 2c-d, Supplemental Fig.2c). Adipose tissue vascularization is crucial determinant of the oxygenation level in the tissue as recently confirmed in a human study (26). Cifarelli *et al.,* showed that adipose VEGFa expression was (a) significantly lower in metabolically unhealthy obese compared to metabolically healthy obese and lean individuals and (b) positively associated with adipose pO2 levels (26). Our genetic model thus provides compelling evidence that deletion of *Phd2* in adipocytes facilitates metabolic protection through multiple effects in functionally distinct adipocyte populations, including enhanced function/plasticity of BAT and enhanced lipid-retention capacity of WAT. Ultimately, this enhances whole animal energy expenditure even at thermoneutrality.

**Figure 2.**
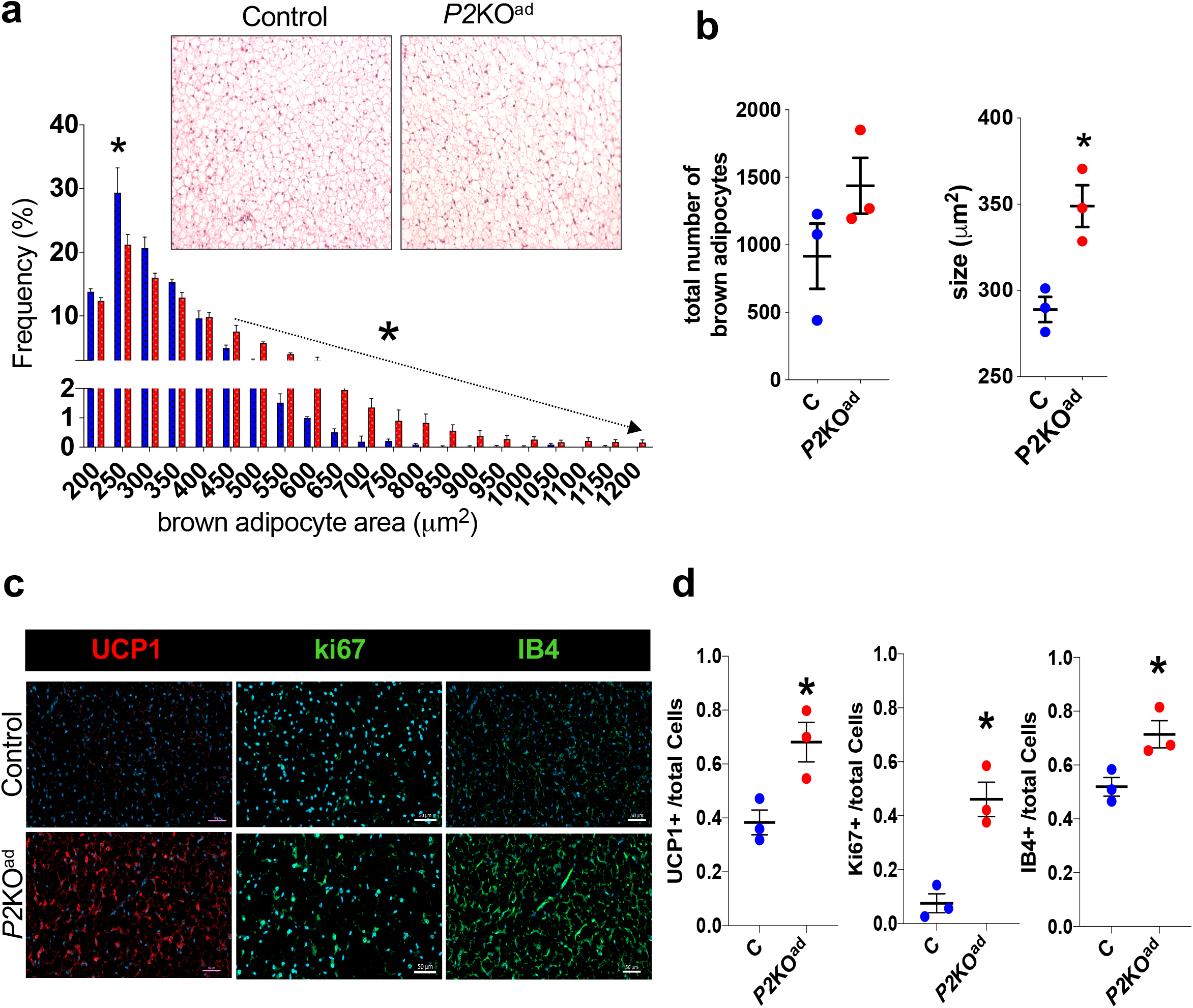
P2KO^ad^ mice maintain functional BAT at thermoneutrality. H&E staining of BAT and frequency distribution of adipocytes **(a).** Quantification graphs of number and size of brown adipocytes in P2KO^ad^ (red) and control littermates (blue) **(b)**. Immunofluorescence images (scale bars 50μm) **(c)** and quantification graphs **(d)** of BAT stained for UCP1+ (red) and ki67+ and IB4+ (green) cells at 28°C (n=3). Nuclei stained with DAPI (blue). Data are presented as mean+/− SEM. *p<0.05, by Student t-test.

Notably, another study, using a whole–body PHD2 inhibition (hypomorphic for PHD2; Hif-p4h-2^gt/gt^), showed reduced body weight, WAT mass and higher EE at room temperature in mice kept on a diet that induced fatty liver and fibrosis (HF-MCD) (27). The authors suggested that improved beiging of WAT could explain their higher heat production (27). WAT beiging may contribute to some degree to increased thermogenesis (28–29), but it is unlikely to be the sole explanation. Pan-tissue PHD2 reduction, as reported by Laitakari *et al.,* will likely affect other tissues with a greater contribution to energy expenditure such as the skeletal muscle or brown adipose. Intriguingly, Hif-p4h-2^gt/gt^ mice exhibited reduced BAT mass (27) in contrast to our adipose-specific *Phd2* deletion mouse model. Although the focus was on NAFLD and liver function (27), the authors did not report effects on skeletal muscle, that could potentially explain enhanced thermogenesis. For example, a recent study showed that global or skeletal muscle deletion of PHD3 led to enhanced exercise endurance capacity and a small increase in maximum oxygen consumption rate (VO2) due to increased fatty acid oxidation in muscle (30). We cannot directly compare the work by Laitakari et al. to our study as they were conducted under different environmental temperatures (RT vs TN), disease (NAFLD vs basal) and genetic models (whole body vs adipose-specific targeting). However, a commonality between the studies is the suggestion that targeting the PHD2 isoform, and specifically in adipose tissue, as clearly shown in our study, may have beneficial lipid lowering effects that could be explored further in humans.

### Pan-PHD inhibition induces UCP1 in vitro

The unexpected finding that PHD2 deletion result in retention of BAT mass at thermoneutral conditions, opens up potential for a new clinically important role of already available oral hypoxia-inducible factor prolyl hydroxylase inhibitors (PHDi) for metabolic disease. The reported PHDis are not isozyme–selective (31), targeting PHD1-2-3 isoforms (pan-PHDi) and are designed for the treatment of CKD (11–13). Phase 3 clinical trials in CKD patients reported that alongside the primary outcomes to increase haemoglobin and erythropoietin levels, at least some PHDi (i.e. roxadustat, dabrodustat) can lower cholesterol and low-density lipoprotein cholesterol levels (32–33). The lowering effects on plasma fatty acids (15) and cholesterol (14,27) have also been shown in animal models treated with PHDi. Targeting BAT activation to alleviate metabolic dysfunction, especially to lower circulating lipid levels, a hallmark of metabolically unhealthy obese (26), is a promising new treatment line, as in both humans and mice activated BAT results in lowering of fasted and postprandial plasma triglyceride levels (34–35). Therefore, it is conceivable that PHD inhibition could be used to target BAT activation. To our knowledge there are no reported studies directly testing whether PHDi enhances brown adipocyte function.

To this end we addressed whether PHDi can confirm induction of UCP1 expression as we have seen in the genetic model of adipose-specific PHD2 deletion. PHDi leads to HIF-1α and HIF-2α stabilization in various experimental models and tissues (15, 36–38) including adipose tissue (15,27). Here, we showed that PHDi increased both protein (Figure 3a) and mRNA (Figure 3b) levels of UCP1 in a mouse immortalized BAT cell line (39). Recent studies showed that the HIF-2a, and not the HIF-1a, signalling pathway was important for BAT function during cold-exposure (40–41), therefore we tested whether the PHD-HIF-2a axis was implicated. Indeed, the effect of PHDi on *Ucp1* and adrenergic receptor *(Adrb3)* mRNA levels was completely abolished (Figure 3b) when brown adipocytes were treated with the HIF-2a antagonist PT-2385 (42), suggesting that PHD2-HIF-2α axis is a crucial regulator of *Ucp1* expression levels. Importantly, PHDi treatment of human abdominal subcutaneous white adipocytes from obese individuals increased *ADRB2* mRNA levels (Figure 3c), a pathway that recently shown to be the main target for pharmacological activation of human brown adipocytes (43). *UCP1* mRNA expression in human adipocytes was also increased by PHDi-treatment (Figure 3d), suggesting that PHDi could induce a *beiging-gene* signature in human WAT.

**Figure 3.**
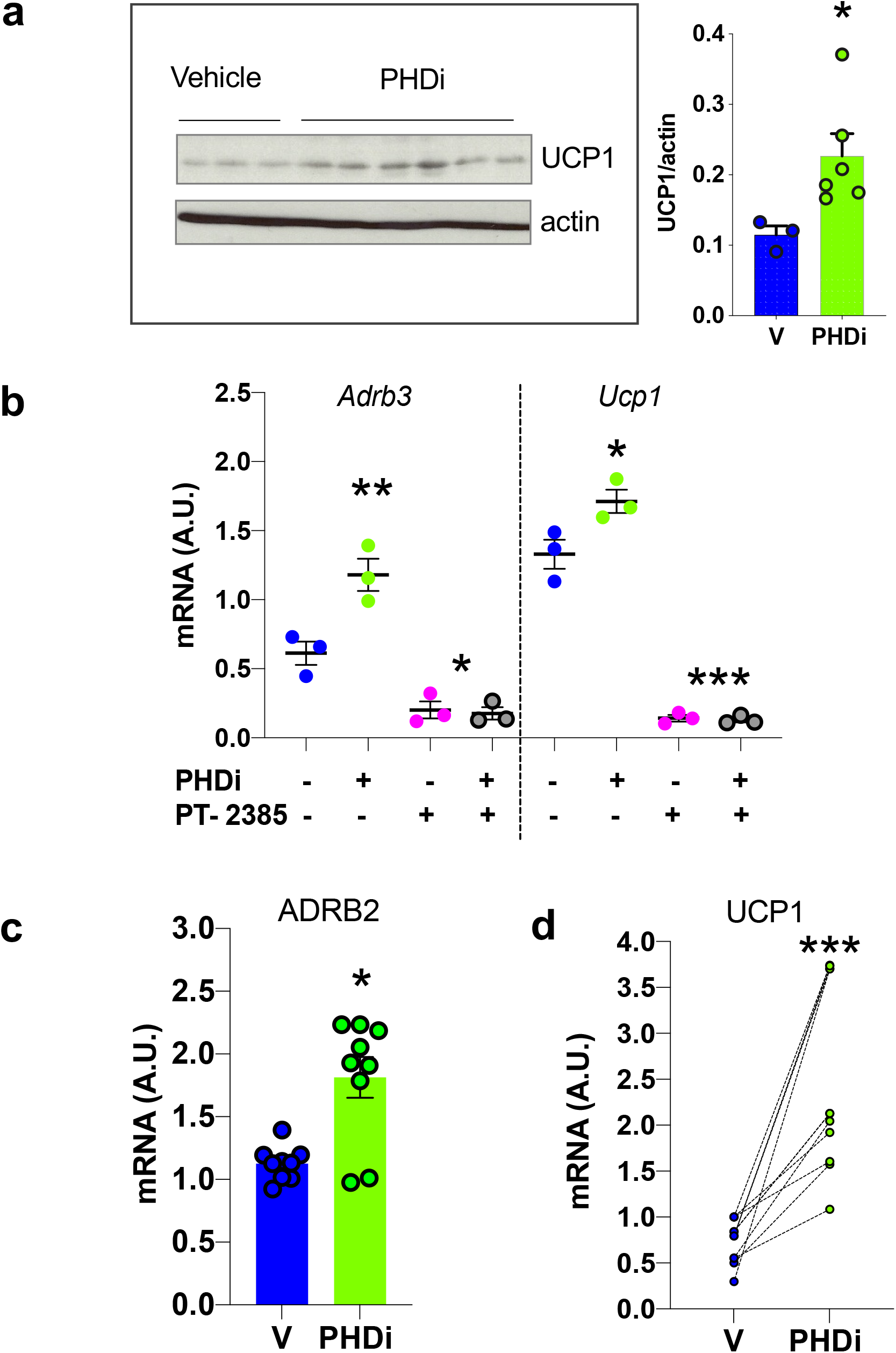
Pharmacological pan-PHD inhibition induces *Ucp1* expression in mouse and human adipocytes *in vitro*. The mouse brown adipocyte cell line (WT-1) treated with PHDi (10μM, 16h, n=6; green circles) showed higher UCP1 levels compared to vehicle (DMSO, n=3; blue circles) **(a)**. mRNA levels of *Adrb3* and *Ucp1* are higher in the PHDi treated (green circles) WT-1 cells. Treatment with the HIF2a antagonist (PT-2385, 10μM, 16h; pink circles) suppressed the effect (n=3) **(b)**. Human adipocytes isolated from abdominal subcutaneous biopsies treated with PHDi (10μM, 16h, n=3 biopsies; 3 replicates per biopsy) show increased ADRB2 **(c)** and UCP1 **(d)** mRNA expression. Data are presented as mean+/− SEM. *p<0.05, ** p<0.01, *** p<0.001 by Student t-test.

### Serum PHD2 protein levels are associated with metabolic dysfunction in humans

Our data here suggest that cellular oxygen sensing proteins, and PHD2 in particular, regulate key metabolic functions and energy homeostasis when targeted in the adipocyte. We next sought genetic evidence for the potential involvement of *PHD2* and downstream hypoxia inducible factor *(HIF* signalling) genes in the human metabolic phenotypes of the ~500,000 UKBioBANK genome–wide association (GWAS) database (using PhenoScanner) (44). Genome–wide significant and/or suggestive association signals were found for blood glucose levels (*PHD2*; rs578226800, p=9.2E^−08^) and basal metabolic rate *(PHD2;* rs7534248, p=3.2E^-06^; *HIF2A;* rs11689011, p=9.5E^-06^; *HIF1AN;* rs1054399, p=6.9E^-10^). Finally, in order to test whether the main human oxygen sensor, PHD2, might serve as a target/biomarker for metabolic disease in humans, we determined if there was any association between serum PHD2 levels and metabolic syndrome traits. We analysed data acquired from a custom version of the SOMAscan proteomic profiling platform screen of serum samples from the large population-based (5457 participants) AGES-Reykjavik cohort (16). The AGES Reykjavik study is a prospective study of deeply phenotyped and genotyped individuals older than 65 years of age. We found that PHD2 levels were significantly positively correlated with visceral adipose tissue (VAT via computed tomography), body mass index (BMI, kg/m^2^), blood markers for type 2 diabetes, HBA1C and insulin, triglycerides and the metabolic syndrome (MetS; Odds Ratio 1.22) (Table 1). The tissue origin of serum PHD2 remains to be determined, however its presence suggests that metabolic dysfunction in ageing is associated with higher PHD2 levels and possibly altered oxygen sensing. This fit with our contention that PHD2 inhibition would have beneficial metabolic effects in humans.

**Table 1.** Human PHD2 serum protein levels are positively correlated with metabolic syndrome related traits. Serum protein PHD2 levels were correlated with BMI, visceral adiposity, triglycerides, fasting glucose, HBA1C, insulin, type II diabetes and metabolic syndrome in the population-based Age, Gene/Environment Susceptibility (AGES) study. N=number of individuals with outcome data, CI=confidence intervals, S.E.=standard error, IFG, impaired fasting glucose.

| Model* | Outcome variable | N | $\beta$ coef. | 95% CI | S.E. | P-value |
| --- | --- | --- | --- | --- | --- | --- |
| **Linear regression** |  |  |  |  |  |  |
|  | Body Mass Index , kg/m<sup>2</sup> | 5439 | 0.356 | (0.234, 0.478) | 0.062 | 1.17E-08 |
|  | Visceral fat area, cm<sup>2</sup> | 5228 | 8.220 | (6.085, 10.355) | 1.089 | 5.20E-14 |
|  | Triglyceride serum, mmol/L | 5447 | 0.066 | (0.048, 0.084) | 0.009 | 6.33E-13 |
|  | Fasting glucose, mmol/L | 5447 | 0.063 | (0.031, 0.095) | 0.016 | 1.05E-04 |
|  | HbA1c, g/dl | 5019 | 0.009 | (0.006, 0.012) | 0.001 | 4.22E-11 |
| | Insulin serum, μU/ml | 5446 | 1.127 | (0.832, 1.422) | 0.150 | 7.36E-14 |
| **Logistic regression** |  |  |  |  |  |  |
|  | Type II diabetes | 5447 | 1.191 | (1.094, 1.297) | 0.052 | 5.76E-05 |
|  | IFG (5.6-6.9 mmol/L) | 4787 | 1.057 | (0.994, 1.124) | 0.033 | 7.53E-02 |
|  | Metabolic Syndrome | 5443 | 1.260 | (1.186, 1.340) | 0.039 | 9.71E-14 |

\*Models were adjusted for PHD2, age and sex. PHD2 beta coefficients for continuous outcome variables are from a linear regression. For dichotomous outcome variables the PHD2 beta coefficients are odds ratios from logistic regression (see Methods for more details on the summary statistics).

In this study we shown that the adipocyte oxygen sensing pathway regulates the thermogenic pathway by sustained UCP1 expression at thermoneutrality. We postulate that this is due to altered plasticity of brown adipose tissue associated with increased angiogenesis and brown adipocyte hypertrophy/hyperplasia that protects against metabolic dysfunction. We further provide evidence for the first time that human serum protein levels of PHD2 are directly correlated with measures of metabolic disease and could potentially be used as a biomarker. In a timely manner, as PHD inhibitors are completing Phase 3 trials or awaiting FDA approval (roxadustat), our findings suggest that this class of drugs, if optimised for isoform and tissue targeting, could be repurposed for treatment of certain metabolic diseases.

## Methods

Further information can be found in the Supplemental Methods and in Supplemental Figures 1-2.

### Human study population

Participants aged 66 through 96 are from the Age, Gene/Environment Susceptibility Reykjavik Study (AGES-RS) cohort (11. AGES-RS is a single-centre prospective population-based study of deeply phenotyped subjects (5764, mean age 75±6 years) and survivors of the 40-year-long prospective Reykjavik study (n~18,000), an epidemiologic study aimed to understand aging in the context of gene/environment interaction by focusing on four biologic systems: vascular, neurocognitive (including sensory), musculoskeletal, and body composition/metabolism. Descriptive statistics of this cohort as well as detailed definition of the various disease relevant phenotypes measured have been published (16–17, Supplemental methods).

### Proteomics data access

The custom-design Novartis SomaScan© is available through a collaboration agreement with the Novartis Institutes for BioMedical Research. Data from the AGES Reykjavik study are available through collaboration ( under a data usage agreement with the IHA.

### Statistics

All data are shown as the mean ± SEM, and a 2-tailed Student’s *t* test was used to compare 2 groups. Data sets were analysed using GraphPad Prism version 8 (San Diego, California). The number of biological replicates is indicated as (n). Unpaired t-test with Welch’s correction was used for the immunofluorescence data. For the inhibitor in vitro experiments, a paired Student’s test was performed to compare effects before and after drug administration in adipocytes from the same individual. A *P* value of less than 0.05 was considered statistically significant.

For the associations of PHD2 to different human phenotypic measures we used linear or logistic regression depending on the outcome being continuous or binary and adjusted for age and sex in our regression analyses. Summary statistics for continuous outcome variables are listed as mean (standard deviation) and median and interquartile range (IQR) for skewed variables. For categorical variables as number and percentages n (%): Body mass index 27.1(4.4), visceral fat area 172.8(80.2), triglycerides 1.0 IQR[0.78,1.43], fasting glucose 5.8(1.2), HbA1c 0.5(0.1), insulin 1.2 IQR[0.79,1.78], type II diabetes 658(12.1%), impaired fasting glucose 1982(41.4%), metabolic syndrome 1677(30.8%).

### Study approval

All animal experiments were conducted according to protocols approved by the U.K. Home Office Animals (Scientific Procedures) Act, 1986. The AGES-RS was approved by the NBC in Iceland (approval number VSN-00-063), and by the National Institute on Aging Intramural Institutional Review Board, and the Data Protection Authority in Iceland.

## Author Contributions

MGS, IPC, RW, KF performed experiments/analysed data. RGM, MB, TC, CJS, RHS provided resources. NMM designed/performed experiments. EFG, LLJ, VG and VE contributed the human proteomics data. ZM conceived the study, performed experiments, supervised research and wrote the manuscript, which was reviewed by all authors.

## Acknowledgements

This work was supported by a British Heart Foundation/University of Edinburgh Centre of Research Excellence Award (RE/13/3/30183 & RE/18/5/34216 to ZM) and a Wellcome Trust ISSF3 (to ZM). CJS and NMM thank the Wellcome Trust for funding. Human adipose tissue collections were supported by a grant from the Chief Scientist Office (SCAF/17/02 to RHS) and the authors acknowledge the financial support of NHS Research Scotland (NRS), through the Edinburgh Clinical Research Facility. The authors are grateful to Professor Sir Peter Ratcliffe (Oxford University) for providing the Phd2^flox/flox^ mice and reviewing the manuscript.

## Conflict of interest

L.L.J. is an employee and stockholder of Novartis. V.E. V.G. C.J.S., N.M.M, R.H.S, M.G.S, T.C., R.G.M., I.P.C, K.F. Z.M declare they have no competing interests.

## References

1. Chen KY et al. Opportunities and challenges in the therapeutic activation of human energy expenditure and thermogenesis to manage obesity. J Biol Chem. 2020;295(7):1926–1942.

2. Marlatt KL, Ravussin E. Brown Adipose Tissue: An Update on Recent Findings. Curr Obes Rep. 2017;4:389–396.

3. Kajimura S, Spiegelman B, Seale P. Brown and Beige Fat: Physiological Roles Beyond Heat Generation. Cell Metab. 2015;22(4):546–559.

4. Overton J.M. Phenotyping small animals as models for the human metabolic syndrome: thermoneutrality matters. Internat J Obes. 2010;34:S53–S58.

5. Speakman JR, Keiher J. Not so hot: Optimal housing temperatures for mice to mimic the thermal environment of humans. Mol Metab. 2013;2(1):5–9.

6. Fischer AW, Cannon B, Nedergaard J. Optimal housing temperatures for mice to mimic the thermal environment of humans: An experimental study. Mol Metab. 2018;7:161–170.

7. Gordon CJ. The mouse thermoregulatory system: Its impact on translating biomedical data to humans. Physiology & Behavior. 2017;179:55–66.

8. Xue Y et al. Hypoxia-independent angiogenesis in adipose tissues during cold acclimation. Cell Metab. 2009;9(1):99–109.

9. Ivan M et al. HIFalpha targeted for VHL-mediated destruction by proline hydroxylation: implications for O2 sensing. Science. 2001;292:464–468.

10. Jaakkola P et al. Targeting of HIF-alpha to the von Hippel-Lindau ubiquitylation complex by O2-regulated prolyl hydroxylation. Science. 2001; 292:468–472.

11. Semenza GL. Pharmacological targeting of hypoxia-inducible factors. Annu Rev Pharmacol Toxicol. 2019;59:379–403.

12. Dhillon S. Roxadustat: First global approval. Drugs. 2019;79(5):563–572.

13. Dhillon S. Daprodustat. First approval. Drugs. 2020;80(14):1491–1497.

14. Rahtu-Korpela L et al. HIF prolyl 4-hydroxylase-2 inhibition improves glucose and lipid metabolism and protects against obesity and metabolic dysfunction. Diabetes. 2014; 63(10):3324–3333.

15. Michailidou Z et al. Adipocyte pseudohypoxia suppresses lipolysis and facilitates benign adipose tissue expansion. Diabetes. 2015; 64(3):733–745.

16. Harris TM et al. Age, Gene/Environment Susceptibility-Reykjavik Study: multidisciplinary applied phenomics. Am J Epidemiol. 2007;165(9):1076–1087.

17. Emilsson V et al. Co-regulatory networks of human serum proteins link genetics to disease. Science. 2018;361(6404):769–773.

18. B. Cannon, J. Nedergaard. Nonshivering thermogenesis and its adequate measurement in metabolic studies. J. Exp. Biol. 2011;214: 242–253.

19. Cui X et al. Thermoneutrality decreases thermogenic program and promotes adiposity in high-fat diet-fed mice. Physiol Rep. 2016;4(10) e12799.

20. Hussain MF, Roesler A, Kozak L. Regulation of adipocyte thermogenesis: mechanisms controlling obesity. FEBS J. 2020;16:3370–3385.

21. Matsuura H et al. Prolyl hydroxylase domain protein 2 plays a critical role in diet-induced obesity and glucose intolerance. Circulation. 2013;127(21):2078–87.

22. Eguchi J et al. Transcriptional control of adipose lipid handling by IRF4. Cell Metab. 2011;13(3):249–259.

23. Lee KY et al. Lessons on Conditional Gene Targeting in Mouse Adipose Tissue. Diabetes. 2013;62(3): 864–874.

24. L J Bukowiecki et al. Proliferation and differentiation of brown adipocytes from interstitial cells during cold acclimation. Am J Physiol. 1986;250:C880–887.

25. Ramseyer VD and Granneman JG. Adrenergic regulation of cellular plasticity in brown, beige/brite and white adipose tissues. Adipocyte. 2016;5(2):119–112.

26. Cifarelli V et al. Decreased adipose tissue oxygenation associates with insulin resistance in individuals with obesity. J Clin Invest. 2020;130(12):6688–6699.

27. Laitakari A et al. HIF-P4H-2 inhibition enhances intestinal fructose metabolism and induces thermogenesis protecting against NAFLD. J of Mol Med. 2020; 98:719–731.

28. Cohen JW, Speigelman BC. Adaptive thermogenesis in adipocytes. Is beige the new brown? Genes Dev. 2013 27(3):234–250.

29. Ikeda K et al. UCP1-independent signalling involving SERCA-2b mediated calcium cycling regulates beige fat thermogenesis and systemic glucose homeostasis. Nat Med. 2017;23:1454–1465

30. Yoon H et al. PHD3 loss promotes exercise capacity and fat oxidation in skeletal muscle. Cell Met. 2020; 32(4):215–228.

31. Yeh TL et al. Molecular and cellular mechanisms of HIF prolyl hydroxylase inhibitors in clinical trials. Chem Sci. 2017;8(11):7651–7668.

32. Chen N et al. Roxadustat for Anemia in Patients with Kidney Disease Not Receiving Dialysis. N Engl J Med. 2019;381(11):1001–1010.

33. Baker KA, Jones JJ. An Emerging Treatment Alternative for Anemia in Chronic Kidney Disease Patients: A Review of Daprodustat. Adv Ther. 2018;35(1):5–11.

34. Bartlet A et al. Brown adipose tissue activity controls triglyceride clearance. Nat. Med. 2011;17:200–205.

35. Berbée JF et al. Brown fat activation reduces hypercholesterolaemia and protects from atherosclerosis development. Nat. Commun. 2015;6: 6356

36. Frost J, Ciulli A, Rocha S. RNA-seq analysis of PHD and VHL inhibitors reveals differences and similarities to the hypoxia response. Wellcome Open Res. 2019;29:4:17.

37. Sugahara M et al. Prolyl hydroxylase domain inhibition protects against metabolic disorders and associated kidney disease in obese, type 2 diabetic mice. J Am Soc Nephrol. 2020;31(3):560–577.

38. del Balzo U et al. Nonclinical Characterization of the Hypoxia-Inducible Factor Prolyl Hydroxylase Inhibitor Roxadustat, a Novel Treatment of Anemia of Chronic Kidney Disease. J Pharmacol Exp Ther. 2020;374 (2) 342–353.

39. Fasshauer M et al. Essential Role of Insulin Receptor Substrate-2 in Insulin Stimulation of Glut4 Translocation and Glucose Uptake in Brown Adipocytes. J Biol Chem. 2000; 275:25494–25501.

40. Garcia-Martin R et al. adipocyte-specific hypoxia inducible factor 2a deficiency exacerbates obesity-induced brown adipose dysfunction and metabolic dysregulation. Mol Cell Biol. 2015;36(3):376–393

41. Krishnan J et al. Dietary obesity-associated Hif1a activation in adipocytes restricts fatty acid oxidation and energy expenditure via suppression of Sirt2-NAD+ system. Genes Dev. 2012;26(3):259–270.

42. Wallace EM et al. A small-molecule antagonist of HIF2a is efficacious in preclinical models of renal cell carcinoma. Cancer Res. 2016;76:5491–5500.

43. Blondin DP et al. Human Brown Adipocyte Thermogenesis Is Driven by β2-AR Stimulation. Cell Metab. 2020;32(2):287–300.

44. Staley JR et al. PhenoScanner: a database of human genotype-phenotype associations. Bioinformatics. 2016;32(20):3207–3209.

