## Supplemental Methods for "Adipocyte-specific deletion of the oxygen-sensor PHD2 sustains elevated energy expenditure at thermoneutrality"

**Animal Studies**. The *Phd2* conditional allele ($PHD2^{f/f}$) (1) on a congenic C57BL/6 background, were crossed with the adiponectin-*Cre* allele (B6;FVB-Tg(Adipoq-cre)1Evdr/J, the Jackson Laboratories) to achieve adipose-specific conditional knockout mice (referred as *P2*$KO^{ad}$). Genotyping PCRs were performed as previously described (1). In all experiments described, control littermates were used for comparisons. Body weight, lean and fat mass were determined by time domain–nuclear magnetic resonance (TD-NMR) (Bruker LF50; http://www.bruker.com) at RT (21$^{o}$C) and after 3d at thermoneutrality (28$^{o}$C; TN). Mice were single housed in calorimetric cages (PhenoMaster, TSE Systems, Chesterfield, MO, USA), at TN for 3days with free access to food and water *ad libitum*, with 12-h light and 12-h dark cycles (7 a.m. to 7 p.m.). Indirect calorimetry was used to measure energy expenditure (W), ($VCO_2/VO_2$), $O_2$ consumption, $CO_2$ production and movement (Counts/hour). Energy expenditure was normalized by lean mass (W/kg).

To pharmacologically inhibit PHD1-2-3, the 2-(1-chloro-4-hydroxyisoquinoline-3-carboxamido) acetic acid (FG-2216), a potent small molecule inhibitor of the PHD enzymes that has been shown to activate HIFα (1). Animals were bred under standard conditions and fed standard chow (product 801151; Special Diet Services, Essex, U.K.) *ad libitum* unless stated otherwise. Male adult mice (16w old) were used in all the experiments. Animal studies were performed under licensed approval in accordance with the U.K. Home Office Animals (Scientific Procedures) Act, 1986. All experiments/measurements were operator/animal handler blinded with data generated by a second individual blinded to treatment until code breaking.

**Blood Parameters:** For non-esterified fatty acids (NEFA), glycerol and glucose measurements, blood was collected after a 4h fast. Glucose concentration was measured by a blood glucose monitoring system (OneTouchUltra2, Lifescan, Milpitas). NEFA (Wako, Richmond VA, USA) and glycerol (Sigma Aldrich, UK) was measured in plasma prepared from blood collected in EDTA-coated microtubes (Sarsted, Leicester, UK).

**Lipolysis assays in adipose explants:** White adipose tissue (inguinal) was dissected after 3d at TN, and 30mg pieces minced and cultured in charcoal-stripped fetal calf serum (FCS)/DMEM phenol-free media for 3hrs. Media was collected on ice and assays performed instantly. Standards and samples of 40 µl media were incubated with WAKO NEFA kit with 100 µl of Reagent 1 (5 minutes, 37 °C) in a clear, flat-bottom plastic 96 well assay plate. 50 µl of Reagent 2 was then added, and the sample incubated for a further 5 minutes. Absorbance of each sample was read at 565 nm, with a background correction at 660 nm. A standard curve (which is linear between 5 μM and 250 μM) was prepared from a 1 mM reference sample (Wako). Glycerol was measured according to manufacturer instructions (Sigma Aldrich, UK).

**Mouse brown adipocyte culture**. The WT—1 cell line has been previously described (2). Briefly preadipocytes were grown in DMEM high glucose (20% FBS). A day after reaching confluence, medium was changed to induction media (Complete DMEM + IBMX 0.5 mM + Insulin 20nM + Dexamathasone 5 μM + Indometacin 125μM + T3 1nM) for 2d. Thereafter, cells were maintained in the differentiation media (Complete DMEM + Insulin 20nM + T3 1nM) for 5-6d. Fully differentiated brown adipocytes were treated with PHDi (FG2216; 10μM) and/or the HIF2α antagonist PT2385 (10μM) for 16h. Adipocytes were lysed with TRIZOL (Invitrogen, Paisley, U.K.) on ice and immediately frozen for further analysis.

**RT-qPCR.** Total RNA was extracted from cells and tissue using TRIZOL (Invitrogen, Paisley, U.K.) and treated with DNase I (Invitrogen, Paisley, U.K.). 1 μg total RNA was used for first-strand DNA synthesis using Superscript III cDNA Synthesis system (Invitrogen, Paisley, U.K.), and qPCR was performed with the Lightcycler 480 (Roche), using mouse-Taqman assays (Life technologies, Paisley, U.K.) for all genes measured. A standard curve was constructed for each gene measured using a serial dilution of cDNA pooled from all samples. Results were normalized to the expression of *Ppia*.

**Immunoblot Assays.** Whole tissue or cell lysates were prepared in ice-cold buffer (5 mmol/L HEPES, 137 mmol/L NaCl, 1 mmol/L $MgCl_2$, 1 mmol/L $CaCl_2$, 10 mmol/L NaF, 2 mmol/L EDTA, 10 mmol/L Na pyrophosphate, 2 mmol/L $Na_3VO_4$, 1% Nonidet P-40, and 10% glycerol) containing protease inhibitors (Complete Mini; Roche Diagnostics Ltd., West Sussex, U.K.). Blots were probed with HIF-1α (1:200; Cayman, Ann Arbor, Michigan), UCP1 (1:000; ab10983, Abcam) antibodies. HRP-conjugated anti-rabbit (Dako, Cambridgeshire, U.K.) secondary antibody was used. Signal was detected using ECL Plus (GE Healthcare Life Sciences, Buckinghamshire, U.K.). Blots were re-probed with an HRP-conjugated anti-actin antibody (abcam, Cambridge, U.K.). Densitometry was performed using the Image J software.

**Histomorphological assessment of brown adipose:** Formalin-fixed, paraffin embedded brown adipose sections (4μm) were used. Images were acquired using a Zeiss microscope (Welwyn Garden City, Hertfordshire, U.K.) equipped with a Kodak DCS330 camera (Eastman Kodak, Rochester, NY). Six randomly selected fields per BAT section in each mouse stained with H&E were captured with 10x objective. Adiposoft software (ImageJ) was used to determine mean adipocyte size ($\mu m^2$) and mean adipocyte count.

**Immunofluorescence in BAT:** Paraffin embedded BAT samples (sections 4μm) were dewaxed by sequential incubations with xylene for 10 min, 100% ethanol for 1 min, 95% ethanol for 1 min, 80% ethanol for 1 min and 70% ethanol for 1 min. After dewaxing, slides went through an antigen retrieval step by boiling the slides in citrate buffer (0.05% Tween, pH 6) for 5 min. After fixation/dewaxing, slides were washed 3 times with PBS for 10 min then blocked with 10% goat serum for 1 h at RT. Sections were then incubated overnight at 4°C with the following primary antibody: UCP1 (1:500, ab10983, Abcam), KI67 (1:50, NB500-170SS, Novus Biologicals) and isolectin B4 (1:200, B-1205, Vector Laboratories). Next day, slides were washed 3 times with PBS for 5 min. Subsequently, sections were incubated for 1 hour at room temperature with species-specific secondary antibodies diluted 1:300. The following fluorochrome-conjugated secondary antibodies were used: anti-rabbit-Alexa 555 IgG, anti-rabbit-Alexa 647 IgG, and streptavidin conjugated 488 (all from Life Technologies). After incubation slides were washed 3 times with PBS for 10 min. Slides were mounted using Fluoramount G (SouthernBiotech, Birmingham, AL) and images were acquired using a fluorescence microscope (Zeiss Observer, Zeiss, Oberkochen, Germany) or a slide scanner (Zeiss, Axio Scan). Images were processed using ZEN Blue lite version (Zeiss). For quantification, Random images were taken from different areas of BAT sections. The images were then split into different fluorescent channels (DAPI, AF488, AF555 or AF647) and analyzed with CellProfiler (3) to measure the median intensity of the proteins of interest (UCP1, IB4, Ki67). Data obtained from CellProfiler was then analyzed with Flowjo software or excel to obtain the number of positive cells. Negative controls images were used to gate the positive population.

**Isolation of mature human adipocytes**. Ethical approval (reference number 15/ES/0094) for the collection, storage and subsequent use of human adipose tissue was granted by The Human Tissue (Scotland) Act 2006 and informed consent was obtained from each participant. Abdominal subcutaneous adipose biopsies were collected from 3 females (48±4 years old) undergoing elective surgery for hernia repair or laparoscopic cholecystectomy in the Royal Infirmary of Edinburgh. Anthropometric characteristics include BMI (37.8±2.5kg/m$^2$), waist to hip ratio (WHR: 0.88±0.023), %fat (43.3±1.8), fat mass (43.5±9kg). Adipocytes were isolated as previously described (1) and exposed to FG2216 (10μM, 16h), and immediately lysed for RNA extraction and RT-qPCR as described above.

**Human study population:** Definition of outcomes used in the present study: body mass index was estimated as kg/m$^2$. Visceral fat area was estimated from computed tomography images at the L4/L5 vertebrae. Area was calculated from single 10mm trans axial images using specialized software. Blood samples were drawn after overnight fasting and centrifuged within 2 hours at room temperature. Triglyceride, fasting glucose and HbA1c were analysed in serum using a Hitachi 912 analyzer (Roche Diagnostics) with reagents from Roche Diagnostics. Serum insulin was measured with a Roche Elecsys 2010 instrument. Type 2 diabetes was defined as fasting serum glucose >7.0 mmol/L or self-reported history of diabetes or the use of insulin or oral glucose-lowering drugs. Impaired fasting glucose was defined in the range 5.6-6.9 mmol/L, excluding individuals with known type II diabetes at baseline. Metabolic syndrome was defined as meeting three of the following criteria: *i)* waist/abdominal circumference >102 cm for men and >88 cm for women, *ii)* triglycerides >1.69 mmol/L, *iii)* high density lipoprotein <1.04 mmol/L for men and <1.30 mmol/L for women*, iv)* fasting glucose >6.1 mmol/L *or* treated diabetes (use of antidiabetic medications - ATC group A10), *v)* systolic blood pressure greater than or equal to 130 *or* diastolic blood pressure greater than or equal to 85 *or* treated hypertension (use of antihypertensive medications).

**Human protein measurements**: an expanded custom version of the SomaScan© platform to include proteins known or predicted to be found in the extracellular milieu was designed (4). PHD2 had its own detection reagent selected from chemically modified DNA libraries, referred to as Slow Off-rate Modified Aptamers (SOMAmers©). To avoid batch or time of processing biases, both sample collection and sample processing for protein measurements were randomized and all samples run as a single set. The 5034 SOMAmers that passed quality control had median intra-assay and inter-assay coefficient of variation, CV < 5%. Hybridization controls were used to correct for systematic variability in detection and calibrator samples of three dilution sets (40%, 1% and 0.005%) were included so that the degree of fluorescence was a quantitative reflection of protein concentration.

(1) Michailidou Z et al. Adipocyte pseudohypoxia suppresses lipolysis and facilitates benign adipose tissue expansion. *Diabetes.* 2015; 64(3):733-745.

(2) Fasshauer M et al. Essential Role of Insulin Receptor Substrate-2 in Insulin Stimulation of Glut4 Translocation and Glucose Uptake in Brown Adipocytes. *J Biol Chem.* 2000; 275:25494-25501.

(3) McQuin C, Goodman A, Chernyshev V, et al. CellProfiler 3.0: Next-generation image processing for biology*. PLoS Biol.* 2018;16(7);e2005970.

(4) Emilsson V et al. Co-regulatory networks of human serum proteins link genetics to disease. Science. 2018;361(6404):769-773.
