## Supplemental Figures for "Adipocyte-specific deletion of the oxygen-sensor PHD2 sustains elevated energy expenditure at thermoneutrality"

#### Supplemental Figure 1

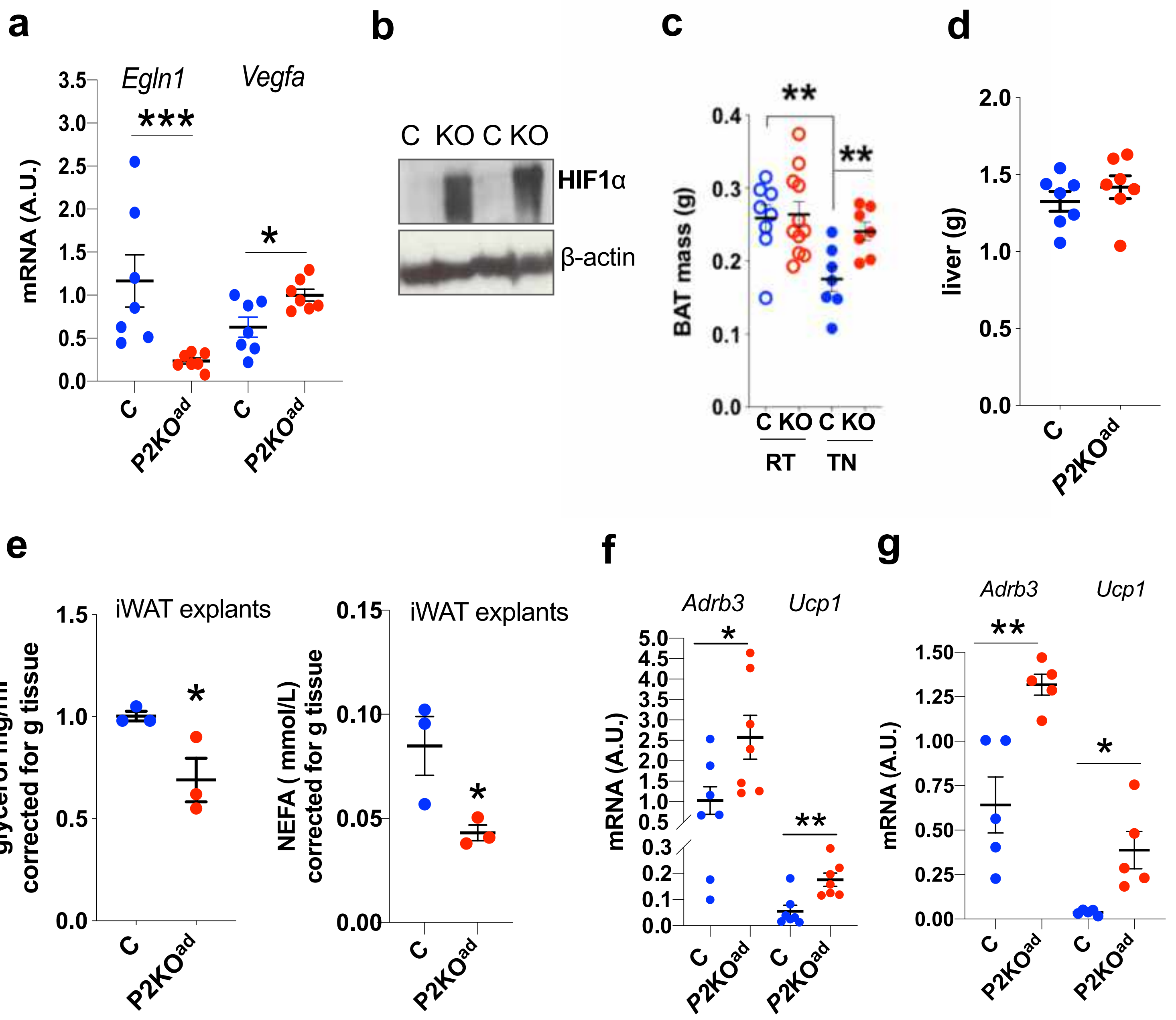


**Supplemental Fig.1. P2KO$^{ad}$ mice retain BAT at TN. (a)** Efficient knock down of *Phd2* (*Egln1*) mRNA in P2KO$^{ad}$ (red) and subsequent induction of the HIF-target gene *Vegfa* (n=7) in BAT. **(b)** P2KO$^{ad}$ mice show stabilization of HIF1α protein in BAT. **(c)** Similar BAT tissue weights at room temperature (21$^{0}$C) in both genotypes (open circles, n=8-11). P2KO$^{ad}$ (KO) mice (red circles) retain but control (C; blue circles) loose BAT mass when housed at TN (28$^{0}$C, n=7). **(d)** Similar liver tissue weight in both genotypes. **(e)** P2KO$^{ad}$ inguinal white adipose (iWAT) explants release less lipids (n=3). **(f-g)** Higher mRNA levels of the beta-3-adrenergic receptor, *Adrb3,* and *Ucp1* in P2KO$^{ad}$ BAT (f; n=7) and iWAT (g; n=5). Significance by Students t-test; *p<0.05, ** p<0.01, ***p<0.001.

### Supplemental Figure 2

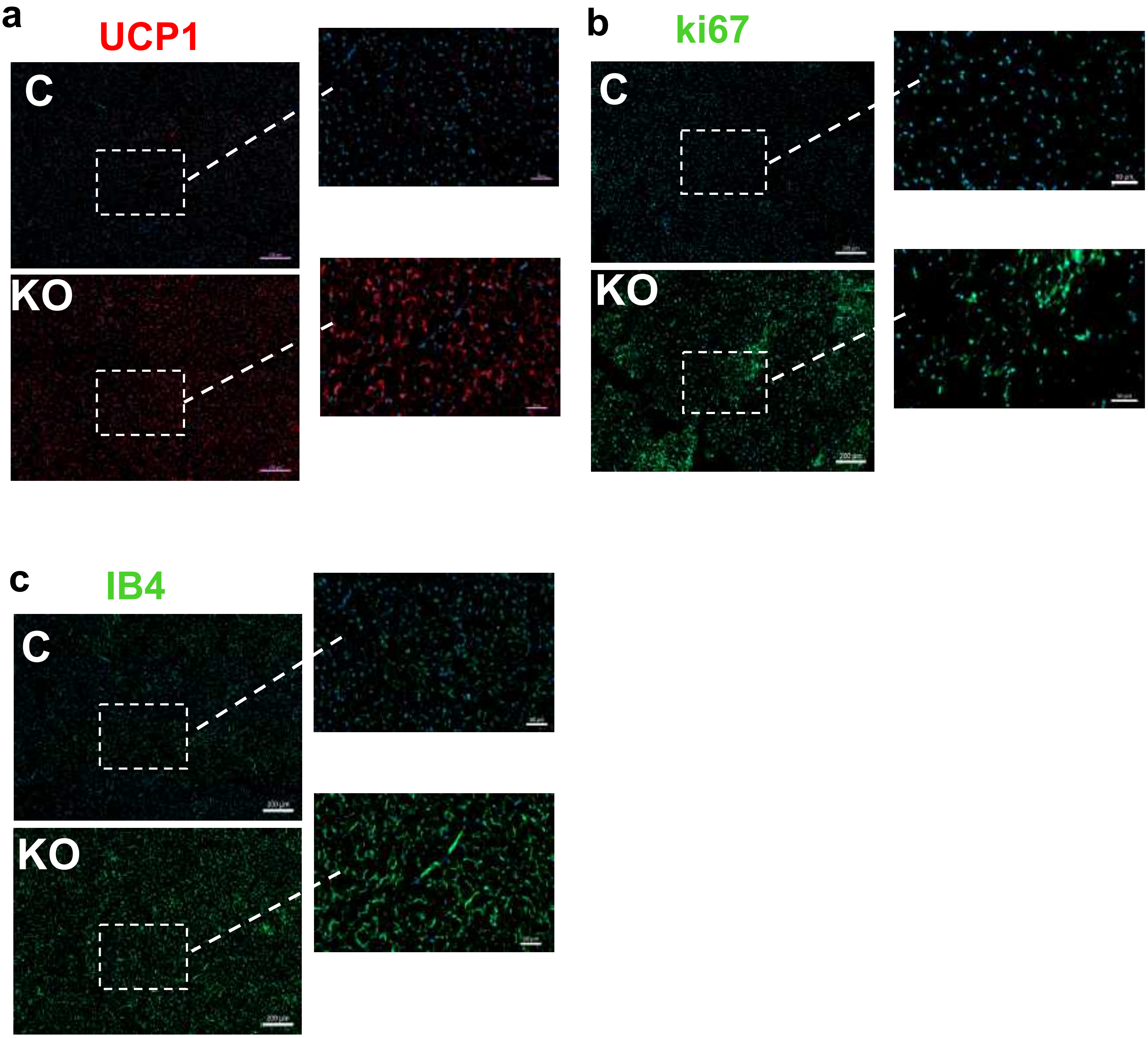


**Supplemental Fig.2. Increased BAT plasticity in P2KO$^{ad}$ mice at TN.** Brown adipose immunofluorescence show higher numbers of UCP1+ cells **(a),** ki67+cells **(b)** and isolectin IB4 staining **(c)** in BAT at 28$^{o}$C (n=3). Images (left) at 200µm and dashed square indicates image selected for higher magnification at 50µm as seen in Figure 2c; C is control, KO is *P2*KO$^{ad}$ .
